# T-cell receptor sequencing of early stage breast cancer tumors identifies altered clonal structure of the T-cell repertoire

**DOI:** 10.1101/172494

**Authors:** John F. Beausang, Amanda J. Wheeler, Natalie H. Chan, Violet R. Hanft, Frederick M. Dirbas, Stefanie S. Jeffrey, Stephen R. Quake

## Abstract

Tumor infiltrating T-cells play an important role in many cancers, and can improve prognosis and yield therapeutic targets. We characterized T-cells infiltrating both breast cancer tumors and the surrounding normal breast tissue to identify T-cells specific to each, as well as their abundance in peripheral blood. Using immune profiling of the T-cell beta chain repertoire in 16 patients with early stage breast cancer, we show that the clonal structure of the tumor is significantly different from adjacent breast tissue, with the tumor containing approximately 3-fold more T-cells, but with a lower fraction of unique sequences and higher clonality compared to normal breast. The clonal structure of T-cells in blood and normal breast is more similar than between blood and tumor and can be used to distinguish tumor from normal breast tissue in 14 of 16 patients. Many T-cells overlap between tissues from the same patient, including approximately 50% of T-cells between tumor and normal breast. Both solid tissues contain high-abundance “enriched” sequences that are absent or of low abundance in the other tissue. Many of these T-cells are either not detected or detected with very low frequency in the blood, suggesting the existence of separate compartments of T-cells in both tumor and normal breast. Enriched T-cell sequences are typically unique to each patient, but there is a subset of sequences that are shared between many different patients. We show that most of these are commonly generated sequences and thus unlikely to play an important role in the tumor microenvironment.

**Significance Statement:** The recent advances in cancer immunotherapy motivated us to investigate the clonal structure of the T cell receptor repertoire in breast tumors, normal breast and blood in the same individuals. We found quantitatively distinct clonal structures in all three tissues, which enabled us to predict whether tissue is normal or tumor solely by comparing the repertoire of the tissue to blood. T cell receptor sequences shared between patients tumors are rare and in general do not appear to be specific to the cancer.

## Introduction

The immune system is thought to play an integral role throughout the life cycle of many cancers including preventing initiation, suppressing development and influencing treatment and patient outcomes(1, 2). Genomic alterations in tumors create immunogenic targets that can be recognized as non-self and eliminated by cytotoxic CD8+ T-cells(3). At the same time this process imposes a selective pressure on the tumor to evade this surveillance(4, 5), sometimes hijacking immune mechanisms that can then be targets for immunotherapy(6, 7). Breast cancer is less immunogenic than other cancers(8), but tumor infiltrating lymphocytes (TILs) have been observed in all subtypes and shown to have prognostic value in triple-negative breast cancer and human epidermal growth factor receptor 2 (HER2)-positive breast cancer [see recent reviews(9-12)].

High-throughput DNA sequencing of the recombined V(D)J region of the T-cell receptor beta chain (TCRB) has become a standard technique for quantifying the distribution of millions of T-cells in a biological sample(13-15). One cell in 100,000 is reliably detected(16), improving clinical monitoring of pathogenic immune cells in various blood cancers(17). Large (>600 individuals) public databases of TCRB data have been generated and used to infer individual MHC alleles and cytomegalovirus (CMV) exposure(18). Even though recent developments in single cell methods can identify both heavy and light chain sequences(19-21), the peptide-MHC target of each T-cell is generally not known. Regardless, characterizing the T-cell receptor repertoire over a range of conditions can provide insight into the subset of T-cell sequences that may be relevant in a variety of clinical applications(22).

Exploratory studies of the T-cell repertoire in tumors from several cancers have found differing repertoires between colorectal tumors and adjacent mucosal tissue(23), intratumoral heterogeneity in renal(24) and oesophageal(25) carcinomas, spatial homogeneity in ovarian cancer(26), two sub-groups of T-cell repertoires in pancreatic cancer(27), increased clonality of CD4+ T-cells in non-small cell lung cancer compare to CD19+ B-cell and CD8+ T-cell compartments(28), and stereotyped shifts in the repertoire after cryoablation and immunotherapy in breast cancer(29). Large scale genomic studies extracting T-cell sequences from bulk RNA-seq data from The Cancer Genome Atlas (TCGA) observed a strong correlation between T-cell diversity and tumor mutation load in a variety of cancer types(30). Exome data from TCGA was used to derive an immune DNA signature for breast cancer that correlated with clinical outcomes(31). Single cell sequencing has also been applied to immune cells infiltrating breast cancer. In one study (20), a subset of CD8+ T-cells with matching alpha and beta chains were detected in breast tumors and sentinel lymph nodes from multiple patients.

In this study we use TCRB sequencing to determine the T-cell repertoire in matched samples of peripheral blood, tumor and adjacent normal breast tissue from 16 patients with early stage breast cancer. We characterize T-cell repertoires from these tissues and show that the clonal structure of the tumor is different from normal breast and blood, and can be used in principle to distinguish tumor from normal. We identify and describe a subset of “enriched” TCRB sequences with high abundance in each tumor and absent or of low abundance in normal breast. Lastly we characterize a subset of CDR3 sequences that are shared between different patients and argue that most of these are likely to be common sequences that are unlikely to play an important role in the tumor microenvironment.

## Results

### Study population

Of 16 invasive breast cancers, 12 tumors were estrogen receptor positive (ER+), progesterone receptor positive (PR+), and human epidermal growth factor receptor 2 negative (HER2-). Among these 12 tumors, 8 were invasive ductal and 4 were invasive lobular carcinomas, grades 1-2 with low proliferation rates (11 of 12 had Ki67 <15%) and 1-3 cm size (10 of 12 less than 3 cm diameter). The remaining 4 tumors had various receptor statuses, including an 8 cm triple-negative (ER-/PR-/HER2-) high-grade tumor with high-proliferation rate (Ki67 of 50-70%) and an ER+/PR+/HER2+ invasive mucinous carcinoma. Two patients received neoadjuvant chemotherapy prior to surgery and one patient had hepatitis C infection (Table 1).

**Table 1.**
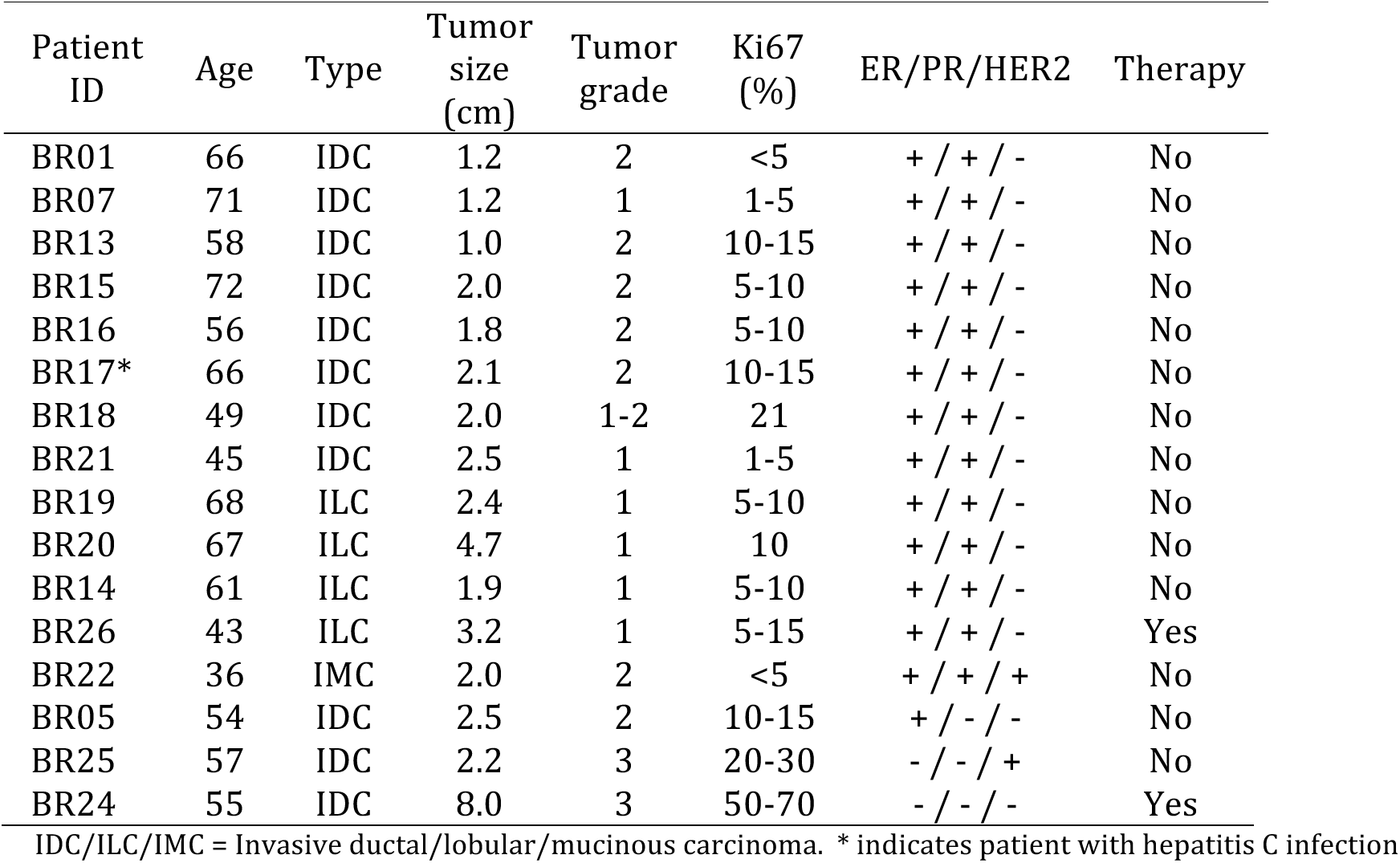
Clinical information for each patient.

| Patient ID | Age | Type | Tumor size (cm) | Tumor grade | Ki67 (%) | ER/PR/HER2 | Therapy |
| --- | --- | --- | --- | --- | --- | --- | --- |
| BR01 | 66 | IDC | 1.2 | 2 | <5 | + / + / - | No |
| BR07 | 71 | IDC | 1.2 | 1 | 1-5 | + / + / - | No |
| BR13 | 58 | IDC | 1.0 | 2 | 10-15 | + / + / - | No |
| BR15 | 72 | IDC | 2.0 | 2 | 5-10 | + / + / - | No |
| BR16 | 56 | IDC | 1.8 | 2 | 5-10 | + / + / - | No |
| BR17* | 66 | IDC | 2.1 | 2 | 10-15 | + / + / - | No |
| BR18 | 49 | IDC | 2.0 | 1-2 | 21 | + / + / - | No |
| BR21 | 45 | IDC | 2.5 | 1 | 1-5 | + / + / - | No |
| BR19 | 68 | ILC | 2.4 | 1 | 5-10 | + / + / - | No |
| BR20 | 67 | ILC | 4.7 | 1 | 10 | + / + / - | No |
| BR14 | 61 | ILC | 1.9 | 1 | 5-10 | + / + / - | No |
| BR26 | 43 | ILC | 3.2 | 1 | 5-15 | + / + / - | Yes |
| BR22 | 36 | IMC | 2.0 | 2 | <5 | + / + / + | No |
| BR05 | 54 | IDC | 2.5 | 2 | 10-15 | + / - / - | No |
| BR25 | 57 | IDC | 2.2 | 3 | 20-30 | - / - / + | No |
| BR24 | 55 | IDC | 8.0 | 3 | 50-70 | - / - / - | Yes |

IDC/ILC/IMC = Invasive ductal/lobular/mucinous carcinoma. \* indicates patient with hepatitis C infection

### Breast Tumors Contain a Larger Proportion of T-cells and a More Clonal Repertoire than Normal Breast Tissue

V(D)J rearrangements within the T-cell receptor beta (TCRB) locus are PCR amplified with V- and J-gene specific primers, and the sequence of the complementarity determining region (CDR) 3 determined using high-throughput sequencing of genomic DNA isolated from peripheral blood, tumor, and normal breast tissue (Fig. 1A). The absolute abundance of input template molecules for each nucleic acid sequence, an estimate of the number of input T-cells, is determined from synthetic spike-in control sequences (see methods section for details). The abundance of individual T-cell clonotypes is determined by combing templates from molecules with the same TCRB V-gene family, J-gene segment and productive (*i.e.*, in-frame) CDR3 amino acid sequence. A total of 2,651,842 templates across 1,097,674 unique clonotypes were detected with individual abundances ranging from 1 to 28,508 templates per clonotype (see Table S1 for details).

**Fig. 1.**
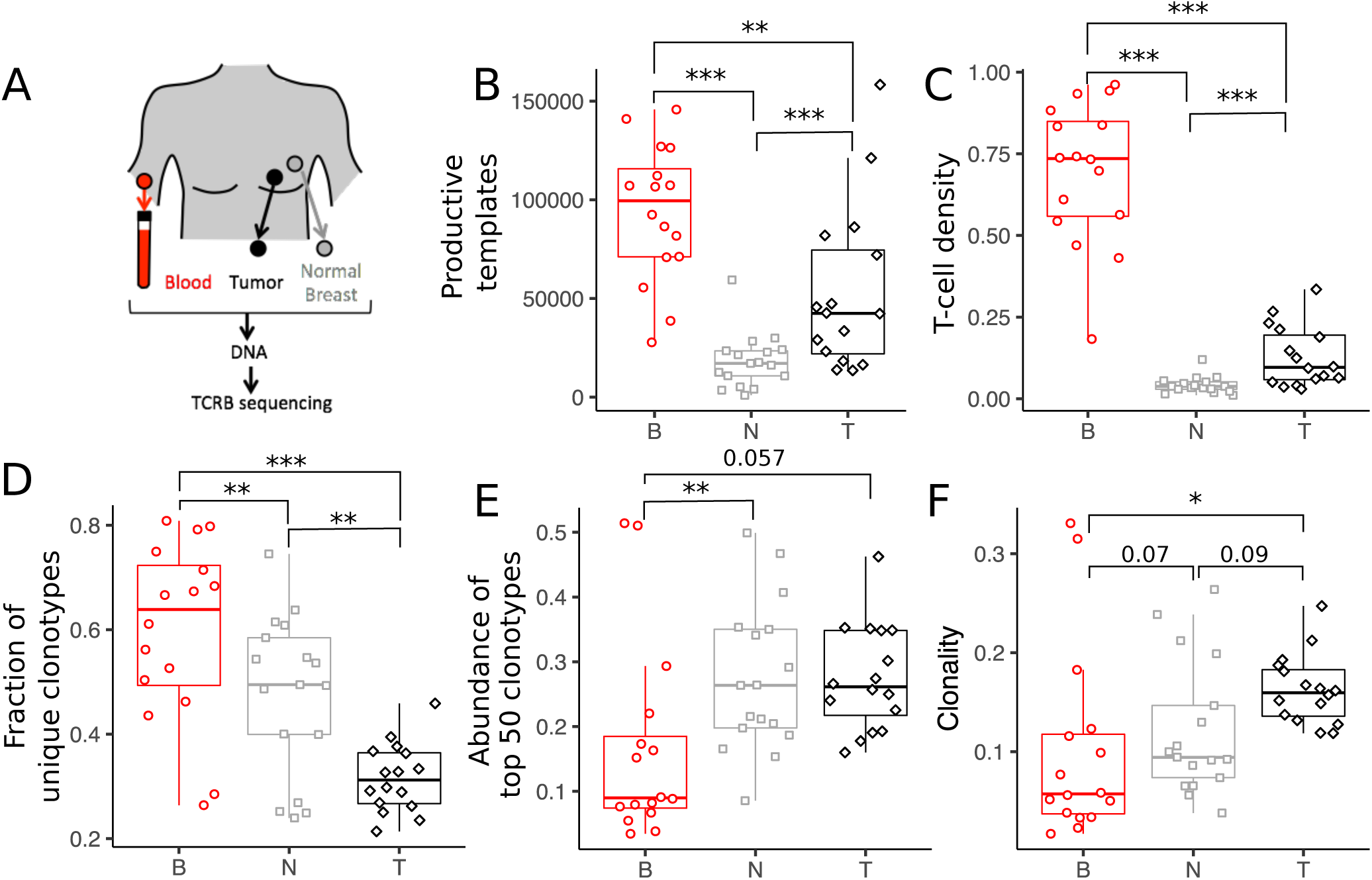
T-cell abundance and repertoire diversity in blood, tumor and adjacent normal breast tissue. (A) Matched trio of samples were obtained from blood PBMCs, tumor and normal breast tissue for each patient. Genomic DNA was isolated from each sample and the T-cell receptor beta chain (TCRB) region PCR-amplified prior to Illumina sequencing. The set of unique clonotypes, defined as TCRB sequences with the same V family, J gene and CDR3 amino acid sequence, and the number of input template molecules for each clonotype comprise the TCRB repertoire in each sample. (B) The number of T-cells in each tissue is estimated from the abundance of in-frame TCRB template molecules, and (C) the number density is determined as the fraction of templates relative to the number of input cells (see Table S1). (D-F) Repertoire diversity for each tissue is estimated from (D) the fraction of templates corresponding to unique clonotypes, (E) the cumulative abundance of the top 50 clonotypes in each sample and (F) the clonality. See Methods for additional details. Unless otherwise indicated, ^*^, ^**^ and ^***^ indicate P-values < 0.05, 0.005, and 0.0005, respectively.

The size of the TCRB repertoire is different for each tissue. The number of productive templates represents the number of T-cells in each sample, and is greatest in blood (median of 99,500 per sample), followed by tumor (42,500) and normal breast tissue (17,200; Fig. 1B). Normalizing the number of productive templates by the number of input cells (see Table S2) results in an estimate of the overall T-cell density in each tissue. Tumors contain approximately 2.5-fold higher density of infiltrating T-cells than adjacent normal breast tissue (median of 9.6% and 3.8%, respectively, P-value <0.0005; Fig. 1C), with both containing a lower density of T-cells than found in circulating PBMCs (74%).

The structure of the repertoires also differs between the three tissues. The number of unique clonotypes is approximately two-fold larger in tumor (median of ~12,500) compared to normal breast (~7,000; P value <0.01) with four-fold more detected in blood (~46,000) compared to tumor (see Fig. S1 and Table S1). The corresponding fraction of T-cells with unique sequences, however, is lower in the tumor (median of 0.31) than in either blood (0.64, P value < 0.0005) or normal breast (0.49, P value <0.005; Fig1E). The cumulative abundance within the top 50 clonotypes is similar between tumor and normal breast (median of ~27%), with both approximately 3-fold higher than blood (9%, Fig. 1D). The clonality in the tumor (median of 0.16) is higher than in blood (0.10, P value< 0.05) and normal breast tissue (0.12, P value <0.09; Fig1F). Both metrics indicate that the repertoire of T-cells in the tumor is less diverse than normal breast tissue, but neither appears to discriminate between ductal (N=8) and lobular (N=4) ER+/PR+/HER2-breast tumors (Fig. S2). The small study size prevents further tumor subtype analysis.

### Abundant Clonotypes Are Often Detected in Multiple Tissues From the Same Patient

Due to the much larger number of sequences in blood compared to either tissue, we calculate the fraction of T-cells in each sample with clonotypes that are detected in all other pairs of samples (Fig. S3). Averaging over the intra-patient tissue combinations shows that approximately half of the templates in tumor and normal overlap with each other and with blood whereas 25-30% of the templates in blood overlap with either tissue (Fig. 2A). Approximately 100-fold less overlap is observed between tissues from different patients (Fig. 2B). Clonotypes with large abundance in one tissue are more frequently detected in other tissues from the same patient (Fig. 2C and Fig. S4). The degree of this overlap is highly variable from patient-to-patient, but the trends are similar within the different tissue compartments of the same patient (Fig. S5). The average overlap across each patient’s tissues is strongly correlated with the average clonality (Fig. 2D) indicating that the observed patient-to-patient variability is likely a property of the underlying repertoire. This is consistent with higher clonality repertoires containing more abundant clonotypes, which have a greater probability of being detected in multiple tissues. Sequences detected in all three tissues are also biased towards high abundance. Considering the trio of samples together for each patient, the fraction of unique sequences detected in more than one tissue varies between 0.5% and 5%, but the fraction of templates detected in all three tissues is nearly 30% compared to less than 10% between any two tissues (Fig. S6).

**Fig. 2.**
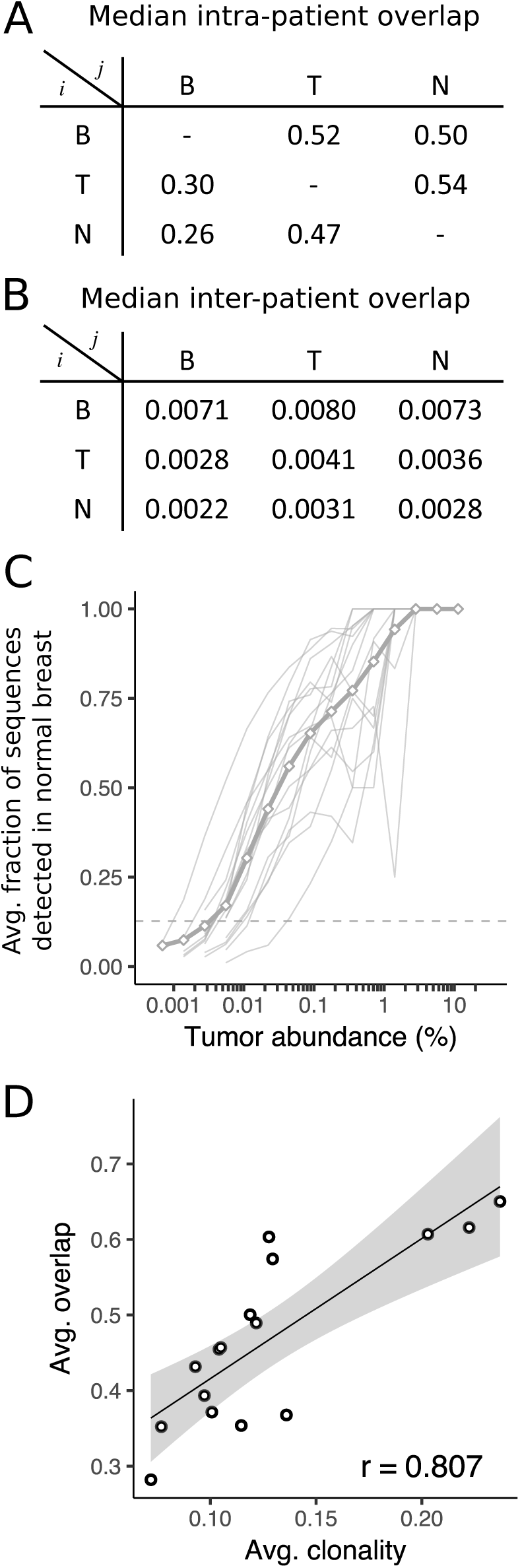
Overlap of TCRB templates between tissues and patients. (A-B) The overlap fraction between two samples is defined as the fraction of templates in tissue *j (columns)* with sequences that are also detected in tissue *i (rows)*, with the median value reported for within (A) and between (B) patient tissue combinations. (C) On average (*thick gray line*) and individually (*thin gray lines*), sequences with high abundance in the tumor are more likely to be detected in normal breast. Similar trends hold for other tissue combinations (Fig. S4). (D) The overlap between samples observed for each patient is strongly correlated (linear regression with 95% confidence interval and Pearson correlation coefficient, *r*) with the clonality of the patient’s repertoire.

### Tumor, Normal Breast, Blood and Contra-Lateral Breast Clonotype Abundances in a Single Patient

Despite the significant fraction of overlapping T-cells between tissues within the same patient, some highly abundant clonotypes in one tissue are either not detected or detected at very low levels in the other tissues. Scatter plots of the abundance of each TCRB sequence between tumor and normal breast for all patients (Fig. S9) indicate that such clonotypes exist over a range of abundances with variable amounts detected in blood (Fig. S10 and Fig. S11).

In patient BR21, for example, 1802 sequences (comprising ~45% of the templates) overlap between tumor and normal and span a broad range of abundances from 0.002% to 11% (Fig. 3A). 154 clonotypes are present with abundance greater than 0.1% in either tissue, including 32 in the tumor (9.5% of templates) that are missing from normal breast and 16 in normal breast (4% of template) that are missing from the tumor. In order to characterize this subset of ‘tumor-enriched’ sequences, we define them to include sequences that are greater than 0.1% abundance in the tumor and at least 32-fold greater abundance relative to normal breast (see Methods for details). In the tumor, approximately half (42 of 81) of the abundant sequences (29% of templates), are enriched whereas in normal approximately one-third (32 of 89) of abundant sequences (9% of templates) are enriched. Unlike BR21, where the top 8 clonotypes in the tumor are all enriched, most patients typically contain several clonotypes with high abundance in both tumor and normal breast tissue (Fig. S9) suggesting that high abundance clonotypes in the tumor are not specific to the tumor.

**Fig. 3.**
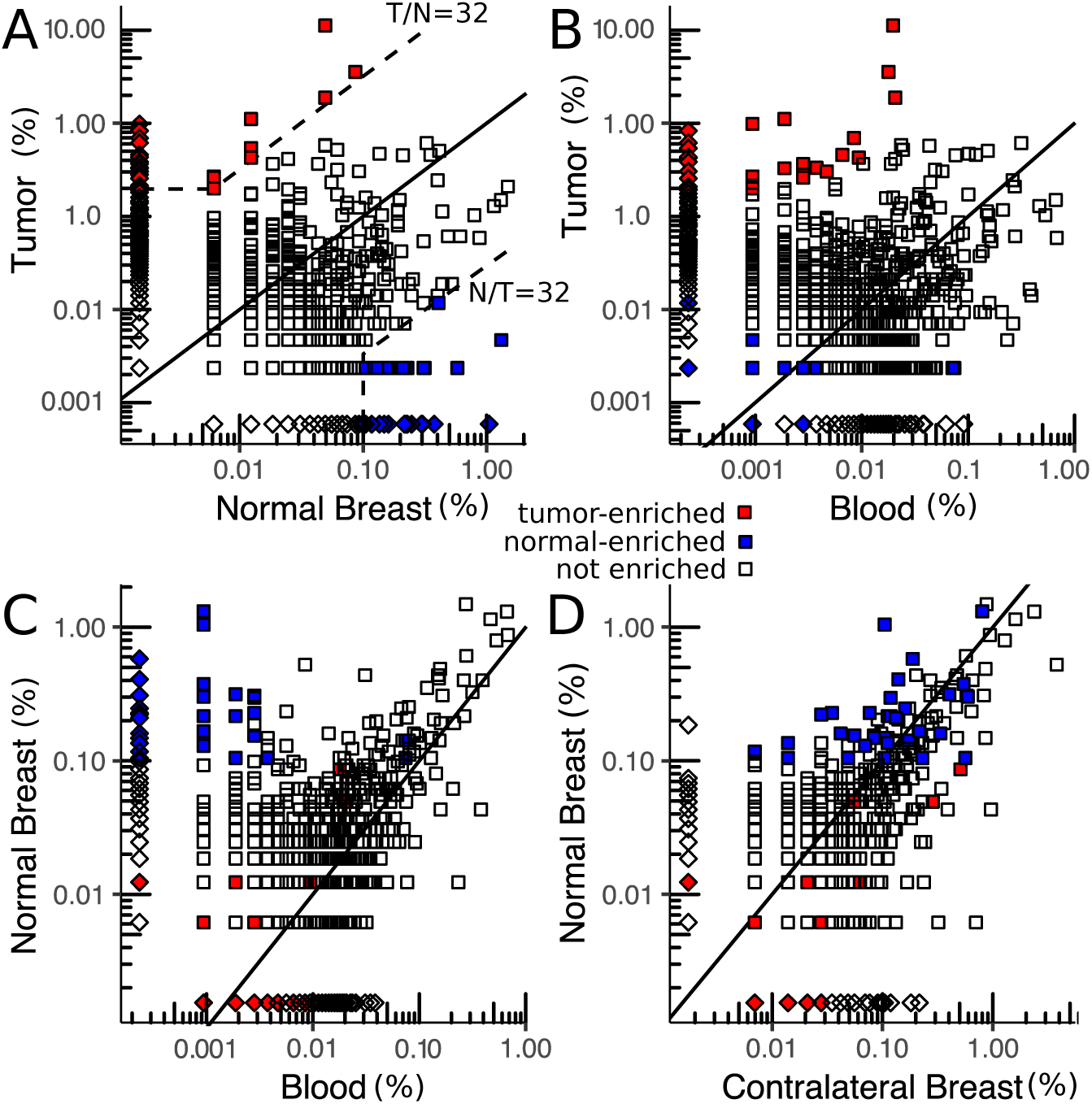
Scatter plots of clonotype abundance for patient BR21. (A) Tumor vs. normal breast. (B) Tumor vs. blood. (C) Normal breast vs. blood. (D) Normal breast vs. contra-lateral normal breast. Each point represents a clonotype detected in one (*diamond*) or both (*squares*) samples. Clonotypes enriched in the tumor relative to normal breast or normal relative to tumor are defined as relative abundance greater than 32 and absolute abundance greater than 0.1% (*dashed lines*). Note that many sequences, especially at low abundance, contain the same number of templates in both samples but are only represented by a single point, see Fig. S7 for the distribution of clonotype abundances.

At the sequencing depth used in this study, approximately 60% of all TCRB templates in tumor and normal breast have clonotypes that are detected in blood (~24% of the templates in blood), including a similar fraction of tumor-enriched (24 of 42; Fig. 3B) and normal-enriched (17 of 32; Fig. 3C) clonotypes. Most tumor-enriched clonotypes correspond to low abundance clonotypes in blood (median is less than 0.001%), which together total only 0.15-0.17% of templates.

BR21 is the only patient with a second normal sample from the contralateral breast (Fig. 3D). The top clonotypes in the two normal tissues are very similar with 109 of 113 sequences (~18% of templates) detected in both tissues. Unlike in blood where the shared clonotypes have very low abundance, the shared clonotypes in the contralateral breast are highly correlated with those in normal breast tissue (Pearson correlation = 0.60, Fig. S8A). All of the normal-enriched sequences are also detected in the contralateral breast.

Pearson correlation coefficients of the abundance from the top 100 clonotypes between each pair of tissues (Fig. S10, Fig. S11) indicate that normal breast, blood and contralateral breast are more correlated with each other (0.45 to 0.6) than with the tumor (-0.13 to -0.03, Fig. S8A). This trend is consistent across many of the patients with larger correlations between normal breast and blood (median correlation of 0.40) compared with normal breast and tumor (0.08) or blood and tumor (0.01, Fig. S8B).

### Classifying Tumor and Normal Breast From the T-cell Repertoire

The differences in clonal structure between tumor and normal (Fig. 1) suggest that the T-cell repertoire could be used to identify a hypothetical biopsy of uncertain tissue as either normal breast or tumor. The larger correlation between normal and blood compared to tumor and blood (Fig. S8B) indicates that including a matched blood sample along with a tissue biopsy would further improve the performance of such an assay. We find that the fraction of the variance in the data captured by a linear model fit to the abundance of the top 100 clonotypes in blood with the corresponding abundance in either tissue (i.e., the coefficient of determination or “R^2^” of the fit lines in Fig. S10 and Fig. S11) distinguishes tumor (median of 0.17) from normal (median of 0.49, P-value < 3.5e5; Fig. 4A). Receiver operator curves (ROC) comparing performance of the R^2^ metric with T-cell density, clonality, and unique T-cell fraction shows that all four have reasonable performance but that the R^2^ metric has a peak sensitivity and specificity of ~0.94 and a larger area under the curve (AUC) of 0.98 compared with the others (AUCs of 0.85, 0.74 and 0.79; see Fig. 4B).

**Fig. 4.**
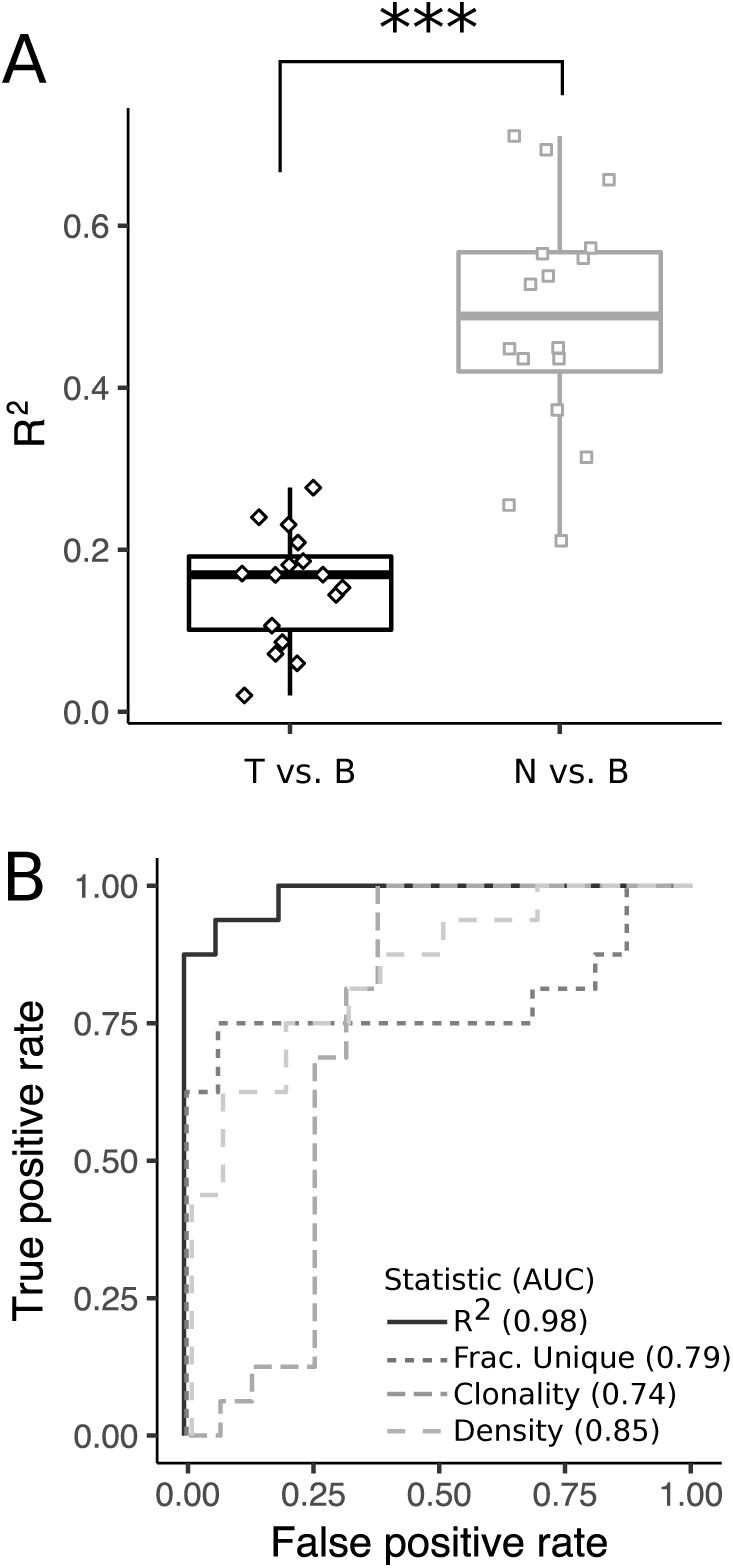
Distinguishing between TCRB repertoires from tumor and normal. (A) The coefficient of determination (R^2^) resulting from fitting the tumor vs. blood abundance and normal vs. blood abundance (see Fig. S10 and Fig. S11 for fit lines). (B) Receiver operator curves and corresponding area under the curve (AUC) values for metrics that distinguish repertoires between tumor and normal breast.

### Subsets of Sequences Are Highly Enriched in Tumors and in Normal Breast Tissues

The number of tumor-enriched sequences varies across patients from 5 to 64 (median of 22) sequences per sample corresponding to 0.9% to 29.4% (median of 6.7%) of templates in each tumor (Fig. 5A). While the exact value of the threshold used to define “enriched” is somewhat arbitrary, the distribution of relative (i.e., Tumor/Normal) abundances indicates that the enriched fraction above ~30 exceeds the amount expected from a log-normal distribution fit to the binned data (Fig. 5C). There also exists a similar distribution of enriched sequences in normal breast tissue where the number of normal-enriched sequences per sample varies between 1 and 65 (median of 18) with template abundances ranging from 0.9% to 29% (median of 6.7%, Fig. 5B). The overall number of normal-enriched sequences also exceeds what is expected from a log-normal fit (Fig. 3D). In contrast, the number of enriched sequences with abundance less than 0.1% in tumor and normal is relatively low and does not exceed the amount predicted from a log-normal fit to the distribution (Fig. S12).

**Fig. 5.**
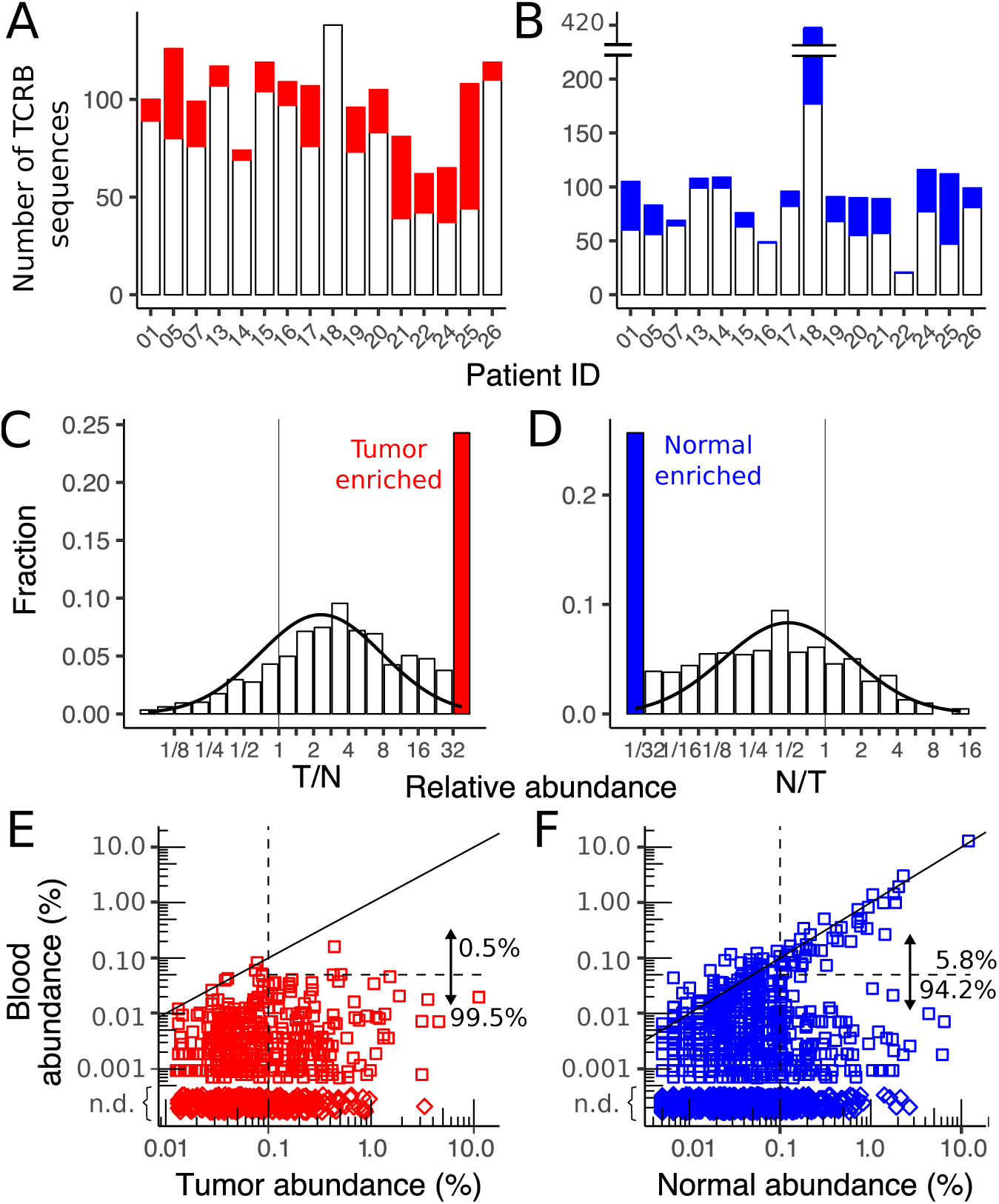
Enriched sequences in tumor and normal. (A-B) The total number of sequences with abundance greater than 0.1% in each patient including the number that are enriched (A) in the tumor relative to normal and (B) in the normal relative to tumor. The relatively large number of enriched sequences in normal breast from BR18 is likely due to low input amount of normal breast DNA in this sample (Table S1), which can also seen in the small number of low abundance (<0.1%) sequences (Fig. S12). (C-D) Distribution of enriched sequences across all patients for sequences in the tumor (C) and normal breast (D, not including the outlier BR18) including a curve fit to a log normal distribution (*solid black line*). (E-F) Scatter plot comparing the enriched clonotypes in tumor (E) and normal breast (F) with their abundance in peripheral blood. Both high and low abundance (0.1%, *vertical dashed line*) clonotypes are shown and the fraction of sequences that are highly correlated with blood (*solid diagonal line*) and with abundance greater than 0.05% (*horizontal dashed line*) enumerated. Clonotypes detected in tumor or normal but not detected (*n.d*.) in blood (*diamonds*) are depicted with arbitrary tissue abundance.

Across all patients, approximately 40% of tumor-enriched and 32% of normal-enriched clonotypes were detected in peripheral blood. For tumor-enriched sequences, there is no correlation between the abundances in tumor and blood with only 2 of 370 tumor-enriched sequence detected above 0.05% abundance in blood (Fig. 3E). In normal, however, 44 of 761 (6%) enriched sequences exceed 0.05% abundance in blood and are highly correlated with the abundance in normal tissue (Fig. 3F). Thus, normal-enriched sequences seem to have two subsets, one with abundances that are correlated with those in blood and the other where they are not. Tumor-enriched clonotypes only contain the subset of sequences that are uncorrelated with the abundance in blood.

### Shared Sequences Between Patients Are Consistent With Common Sequences

The subset of TCRB CDR3 sequences shared across multiple samples may contain clonotypes that recognize a common shared antigen, such as a breast cancer epitope, or to sequences that occur naturally across multiple people. We show below that commonly shared CDR3 sequences have similar properties in the three tissue compartments and, unlike tumor-enriched sequences, are consistent with commonly occurring CDR3 sequences in a population of healthy donors.

The fraction of clonotypes with shared CDR3 sequences between two patients across the different tissue combinations are well described (within ~20%) by a standard capture-recapture model with a single fit parameter representing the total number of productive sequences in the population (*M*=2×l0^6^,Fig. S13). In order to compare the degree of sharing between different tissues, we compare the overlap amongst the top ~2000 most abundant CDR3 sequences in each tissue (see Methods). Using the inferred population size M, we simulated the expected fraction of sequences detected in multiple patients and show that the amount of sharing exceeds what is expected from random sampling (Fig. 6A). Approximately 2% of abundant tumor and normal breast sequences and 4% of those in blood are shared across two or more patients. In particular, the tail of the distribution is larger than expected with 102, 13 and 6 highly shared sequences detected in more than 5 patients from blood, tumor and normal breast, respectively. The capture-recapture model ignores clonotype abundance within each patient, but this assumption is consistent with the minimal bias observed in the abundance of highly shared CDR3 sequences compared with unshared sequences (Fig. 6B).

**Fig. 6.**
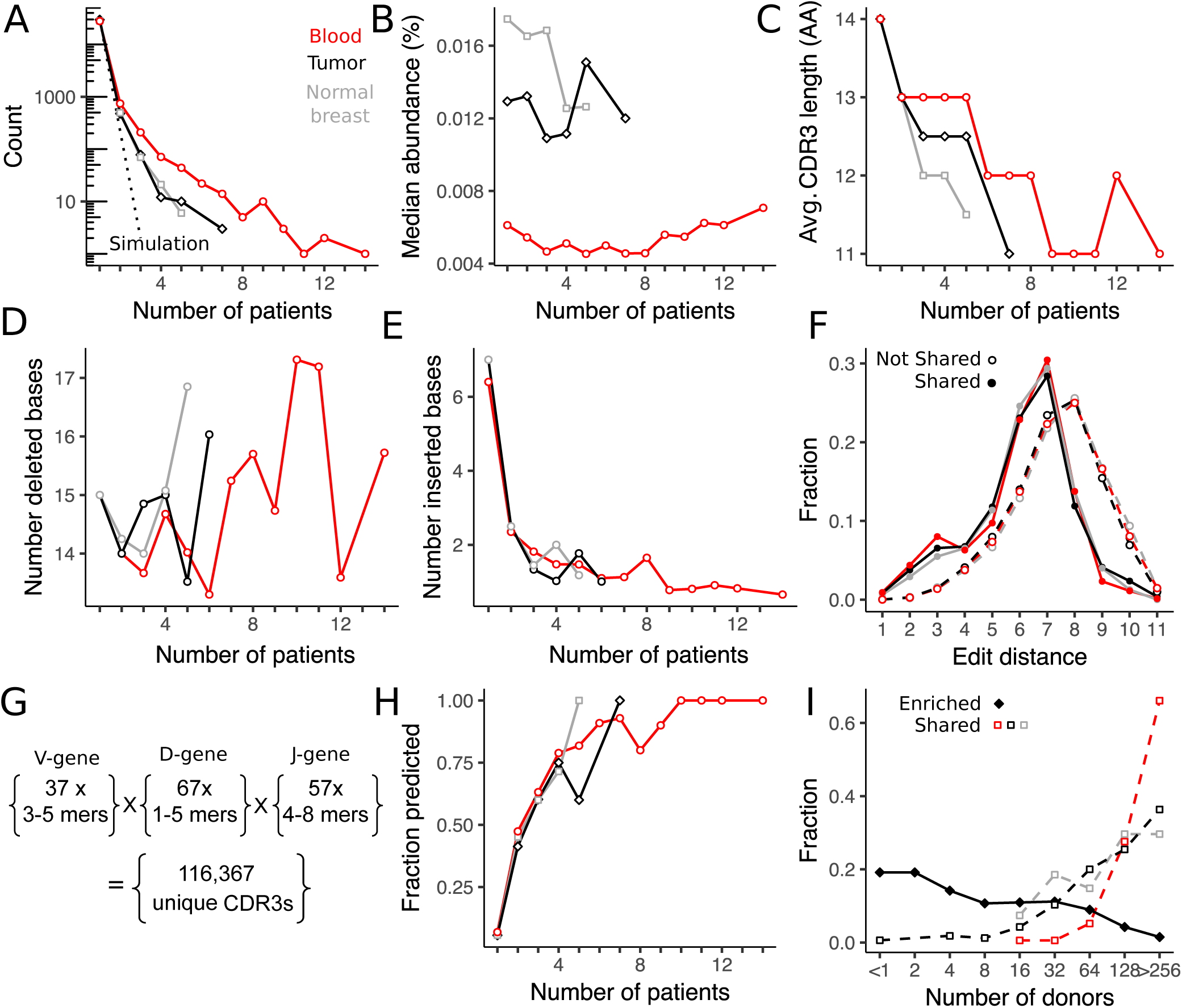
Inter-patient T-cell sharing of clonotypes between tissues. (A) The fraction of blood, tumor, and normal breast tissue sequences found in one (not shared) or more (shared) patients compared to the expected fraction by random sampling (*dashed lines*). (B) The median abundance of clonotypes detected in multiple patients is not sensitive to the degree of sharing. (C-E) Compared with clonotypes that are not shared across multiple patients, shared clonotypes have (C) shorter CDR3 amino acid length, which can be attributed to a similar number of (D) deleted but fewer (E) inserted nucleotide bases in the junctional regions between V-D and D-J genes. (F) The diversity of CDR3 sequences is estimated from the distribution of edit distances (length = 13 amino acids) which is shifted to lower values for shared sequences. (G) The number and length of germline V-, D- and J-gene amino acid sequence fragments used in a recombination model of low diversity CDR3 sequences, and (H) the fraction of CDR3 sequences detected by the model, see Methods for details. (I) Results from a database query of 585 healthy donors (18) indicates that shared CDR3 sequences are also likely to be shared in a population of healthy donors whereas enriched CDR3 sequences are much less likely to be shared in the general population.

Several metrics indicate that highly shared sequences are less diverse than private sequences. First, average CDR3 length is shorter for shared sequences, decreasing from 14.5 amino acids in non-shared sequences to ~11 amino acids in the most highly shared sequences (Fig. 6C). The number of nucleic acid bases deleted from the CDR3 region of shared sequences is relatively unchanged (Fig. 6D), but the number of inserted bases in the CDR3 region decreases sharply from 6-7 nucleic acids in non-shared sequences to ~1 nucleic acid in shared sequences (Fig. 6E). Lastly, the edit distance between all pairs of randomly selected non-shared sequences with length 13 amino acids has a relatively smooth distribution that is peaked at 8 amino acid bases. In contrast, the edit distance distribution between shared CDR3 sequences is shifted toward shorter lengths with a shoulder at approximately 3 amino acid bases indicating a less diverse subset (Fig. 6F).

We used a simple model to compute low-diversity recombinations from all germline amino acid contributions from the V, D and J gene segments to the CDR3 region. The artificial CDR3s include amino acid deletions but no insertions and results in 116,367 unique sequences (Fig. 6G and Methods for further details). 80% of these are not detected in our study, mostly due to non-uniform V and J gene usage in actual repertoires, 10% were detected in non-shared clonotypes with the remaining 10% distributed across all shared CDR3s. After sub-sampling, the fraction of shared CDR3s predicted by the model increases rapidly with the degree of sharing with 100% of the most frequently observed clonotypes in each tissue predicted by the model (Fig. 6H).

Lastly, shared CDR3 sequences in tumor, blood and normal breast are almost all relatively common in a database of T-cell repertoires from 585 healthy donors(18) (Fig. 6I). In contrast, tumor-enriched sequences are much less frequently observed in the database with ~50% of CDR3s detected in less than 5 donors. Both tumor-enriched and shared CDR3 sequences are similarly represented in male and female donors (Fig. S15), which is expected for the commonly shared sequences but not enriched sequences, suggesting that the most highly enriched T-cells in the tumor may not be recognizing antigens specific to female breast cancer.

## Discussion

In this work we performed bulk T-cell repertoire sequencing from peripheral blood, adjacent normal breast and tumor from 16 prospectively collected breast cancer patients, most of which were early stage, treatment naïve, ER+/PR+/HER2-subtype tumors. Advantages of bulk repertoire sequencing include its high sensitivity for detecting T-cell clonotypes(16), which may be missed by traditional staining(29), and the simplicity of genomic DNA input, which is not subjected to the variability of single cell isolation techniques(32) required in cytometry. The disadvantage is that some important cellular details are missed such as the heavy and light chain pairing and the expression of functional markers such as CD4, CD8, FOXP3, CTLA4 and PDL1(19, 33), unless cells have been previously sorted(15, 34). Our goal was to characterize the differences in repertoires between adjacent normal breast tissue and tumor in order to determine if there was evidence for subsets of T-cells that might play a role in the tumor microenvironment(35).

Recent guidelines have been established for evaluating TILs in breast cancer via standard hematoxylin and eosin (H&E) stain(36) and clinical studies have shown that outcomes are correlated with the number of infiltrating T-cells, especially for triple-negative and HER2+ subtypes(37). Quantifying T-cells via sequencing is highly reproducible and correlates well with H&E staining but with higher sensitivity(29). Immunohistochemistry indicates that T-cells in healthy breast tissue are mostly located in lobules with CD8+ T-cells approximately 10-fold more abundant than CD4+(38). Immune infiltrates isolated from tumor and normal breast contain similar amounts of CD8+ T-cells (~20% of T-cells) whereas CD4+ T-cells in tumor comprise ~40% of T-cells compared to ~20% in normal breast tissue(39). We show that repertoires from blood PMBCs, tumor and normal breast are distinct with large differences in the number and density of T-cells detected in each tissue compartment. The tumor repertoire is less diverse than in normal breast as indicated by the lower fraction of unique T-cells and increased clonality in the tumor (Fig. 1E-F). These results are consistent with a similar study comparing the effects of cryoablation and immunotherapy on TCRB repertoires in early stage breast cancer patients(29).

These repertoire metrics have predictive value and could be used to classify unknown tissue sample as taken from tumor or normal breast. The performance in this small cohort varies from an AUC of 0.74 for clonality to 0.85 for T-cell density. Since repertoires from normal breast and blood are more similar than tumor and blood, a better metric can be determined by comparing the tissue and blood repertoires. Using the R^2^ goodness-of-fit between each tissue with blood, the AUC increases to 0.98 with only two samples not properly classified.

Extensive overlap between tissues within the same patient is expected due to normal blood perfusion. On average ~50% of T-cells in either tissue sample were found to overlap with the other tissue or blood samples from the same patient. This fraction varied from patient-to-patient and was correlated with the average clonality of the repertoires in each patient, which is consistent with the more clonal repertoires containing larger clonotypes that are more frequently sampled in different tissues.

In addition to many frequently overlapping clonotypes, most patient tumors contained a subset of clonotypes (median number of 22) that are highly enriched in the tumor relative to normal breast. In a few patients (BR05, BR21 and BR22 in Fig. S9) these clonotypes are clearly dominant in the tumor, however in most patients high abundance clonotypes in the tumor are also highly abundant in the normal suggesting that they are less likely to play an important role in the tumor micro-environment. Methods have been developed to identify the subset of clonotypes that change abundance in serial blood samples following an external perturbation(40). Analogous subsets are not as clear between breast tumors and normal tissue but there is a population of highly enriched clonotypes in each tissue across the cohort of patients (Fig 4B,D). We focused our analysis of enriched clonotypes to those with at least 0.1% abundance due to limited sequencing depth and multiple reports that tumor-reactive T-cells identified in other cancers typically exceed 0.1%. For example, in melanoma and early stage non-small cell lung cancer (NCSLC), ~3%(41) and 0.2-1.5%(42) of infiltrating CD8+ T-cells are tumor reactive with sorting on PD1+ in melanoma further enriching the population to 1-10%(43). CD4+ T-cells may also be tumor-reactive(44, 45) with 0.13-0.3% of expanded CD4+ T-cells tumor reactive in melanoma(46).

Enriched clonotypes in normal breast tissue show two subsets in peripheral blood, one that is highly correlated with blood and the other distributed over a range of low (or zero) abundance in blood. We interpret the first subset as sequences from blood that perfuse the normal tissue whereas the second subset may represent a compartment of tissue-resident T-cells in normal breast(47). The abundance distribution of tumor-enriched clonotypes in blood resemble this second subset seen in normal breast and do not contain clonotypes that are highly correlated with blood. While some of the tumor-enriched T-cells may play a direct role in the tumor microenvironment, many T-cells in this population may instead represent a compartment of tissue-specific resident T-cells that are distinct from blood and performing standard immune surveillance(48).

Public T-cells expressing the same TCR sequence but generated in different individuals have been identified for numerous infectious diseases, autoimmune disorders, and some cancers(49). In cancer, this process is thought to require T-cells that recognize epitopes from mutated proteins that are presented by similar major histocompatibility (MHC) molecules(3), and thus occur more often in cancers with larger numbers of mutations(8). We detect a small fraction of TCRB CDR3s across multiple tumors, but our analysis suggests most of these are low-diversity sequences that are unlikely to be tumor-specific. First, CDR3 sequences shared across tumors follow similar trends as those shared in blood and normal breast, with a similar amount detected in normal breast as in tumor. Shared clonotypes were not strongly biased by abundance and contained shorter CDR3 lengths with many fewer inserted nucleotide bases into the junction regions, as reported in earlier studies(15, 50). A simple model generating all CDR3 recombinations from germ-line amino acid sequences with minimal insertions was able to predict many of the shared sequences including all of the most highly shared sequences. These findings are consistent with more detailed physical models of the nucleotide sequence recombination(51) that show commonly occurring sequences have a higher likelihood of being generated.

Shared and tumor-enriched sequences are distinct, with only two CDR3s detected in both sets. We also queried a large public database of T-cell repertoires in blood from 585 healthy volunteers(18). Enriched sequences were detected in many fewer healthy donors than shared sequences, supporting the hypothesis that enriched sequences may play a specialized role in a subset of patients whereas most shared sequences are common in the population. Interestingly, tumor-enriched sequences were equally prevalent in male and female volunteers, suggesting that they may not be specific to breast cancer.

## Conclusions

By comparing bulk T-cell repertoires from tumor and normal breast tissue in a cohort of early stage breast cancer patients we were able to determine the clonal structure of infiltrating lymphocytes. These clonal structures were quite distinct between tumor and normal and we were able to distinguish tissues as normal or tumor solely on the basis of comparing T-cell repertoire with blood. We identified a subset of T-cells in each patient that were highly abundant in the tumor compared to normal breast; this population may represent a clinically relevant subset for future investigation. In many patients, however, these clonotypes were dominated by more abundant clonotypes, which were also highly abundant in normal breast, thus highlighting the large amount of background T-cells in the tumor that complicate isolating and identifying tumor-specific T-cells.

## Materials and Methods

### Sample Collection

Informed consent was obtained from patients with invasive breast cancers greater than one cm under a Stanford IRB-approved research protocol #5630. 5-10 ml of whole blood was collected in EDTA tubes prior to surgery and processed within 2 hours. After tumor resection, tumor tissue was visually identified, and a portion was excised and placed in prechilled RNAlater. Samples of adjacent normal breast tissue were obtained from sites 2-4 cm away from the tumor, rinsed with PBS and placed in a separate tube of pre-chilled RNAlater. For one patient undergoing simultaneous double mastectomy, normal breast tissue was also obtained from the contralateral breast.

### Sample Processing

Plasma was removed after separation by centrifugation for 10 min at 1600 RCF. The remaining blood cells were re-suspended in PBS and mononuclear cells were isolated via Ficoll-Paque centrifugation with Sepmate-50 tubes (Stem Cell Technologies, Vancouver, Canada). Cells were cryopreserved in 90% fetal bovine serum and 10% DMSO, divided into aliquots containing 2-4 million cells and stored in a liquid nitrogen until use. Cells were thawed at room temperature, diluted with PBS, pelleted at 350 RCF and resuspendened in RLTplus buffer (Qiagen) with 1% betamercaptoethanol (Sigma) and 0.05% Reagent DX anti-foaming agent (#19088, Qiagen). Lysate was passed through a QIAshredder (Qiagen) and DNA and RNA were extracted using an AllPrep DNA/RNA Mini kit (Qiagen) following manufacturer’s instructions.

### DNA isolation

Tumor and normal breast tissue samples were divided into pieces, incubated in RNAlater overnight and transferred to -20°C the following day. For the tumor, 20-50 mg of tissue was diced with a fresh razor blade and homogenized in RLTplus buffer containing 1% beta-mercaptoethanol and 0.5% reagent DX using a Tissuelyzer II (Qiagen) at 30/s for 3-9 min. Homogenized lysate was centrifuged at 20,000 RCF for 3 min, the supernatant removed and passed through a QIAshredder before proceeding with Allprep Mini kit for DNA and RNA isolation. An on-column digestion with 20 ul of proteinase-K (Qiagen) dissolved in 60 ul of Buffer AW1 was implemented prior to washing the DNA column with buffer AW2. DNA yield from tumors was variable but typically 20-50 mg of tumor was required for >3 ug DNA. Normal breast tissue was processed similarly except 100-300 mg of tissue was often required to obtain sufficient (>3 ug) DNA due to high lipid content and low density of cells. In particular, care was taken to avoid the lipid layer after the 20,000 RCF centrifugation step. DNA was quantified with fluorometric quantitation (Qubit High Sensitivity DNA kit) and UV spectroscopy (Nanodrop 1000). In order to minimize any contamination, samples were processed at separate times and all work performed in a PCR cabinet (AirClean).

### Immunosequencing

T-cell receptor beta chain sequencing was performed on genomic DNA purified from blood, tumor and normal breast tissue using the ImmunoSeq Intro kit (Adaptive Biotechnologies) following manufacturers instructions. Briefly, two gDNA replicates per sample, each containing either 0.5 μg (blood) or 1.5 μg (tumor, normal, see Table S 1 for details) are independently amplified using 2× Multiplex PCR Master mix (31 cycles; Qiagen #206151) with proprietary primers and a spike-in control sequences for absolute quantification(13),(52). The product is cleaned up with SPRI beads, and amplified again with 8 additional cycles of PCR to attach Illumina sequencing adapters containing sample-specific barcodes. An 87-bp fragment including the CDR3 region and flanking V and J genes is sequenced using single-ended 150 bp reads of 7 samples (14 replicates) multiplexed onto a single MiSeq lane and sequenced using version 3.0 chemistry. The raw sequencing data is uploaded to Adaptive Biotechnologies and processed using their proprietary pipeline. Results for each sample are reported to the user after merging data from both replicates and includes absolute quantification of template molecules for all nucleic acid sequences detected; CDR3 amino acid sequence for in-frame molecules; V-, D- and J- gene segment identification; the most likely number of bases deleted from each gene segment contributing to the CDR3 and the most likely number of bases inserted into either junction. The average read coverage of each sample was ~10X (see Table S 2 for details).

## Data Analysis

Processed sequence data for each sample was downloaded from Adaptive Biotechnologies and analyzed with custom scripts in bash, awk, and R. Unless otherwise noted, clonotypes are defined here as productive recombinations containing a V-gene family, CDR3 amino acid sequence and J-gene segment. Sequence abundance is calculated as the number of templates for that sequence divided by the total number of templates in the sample. The T-cell density is determined by dividing the total number of templates by the total number of input cells estimated from the input amount of DNA. Clonality is defined as one minus the normalized Shannon entropy of the TCRB abundances (*i.e.*, the Pielou evenness, see Table S1). P-values between tissues are calculated as a paired Wilcoxon rank sum test in R unless otherwise noted.

### Overlapping templates

The fraction 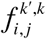 of templates in patient *k*, tissue *j* that are shared with patient *k*’, tissue *i* is defined as the sum of the abundances of all sequences in sample (*j*,*k*) that were also detected at least once in (*i*,*k*’). As a result, 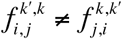 and the fraction of overlapping sequences between tissues with large differences in total numbers of sequences can be represented (Fig. S3). Intra- and inter-patient averages are determined by finding the median over tissue pairs (*i*,*j*) when *k*=*k*’ and *k≠k*’, respectively (Fig. 2A-B).

### Capture-recapture model

The number of overlapping sequences 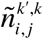 between tissues *(i*,*j)* are modeled as

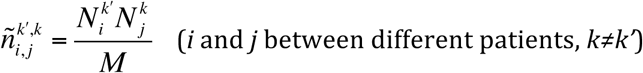

where 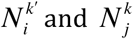 are the unique sequences observed in each tissue, and *M* represents the total number of unique sequences in the study population and is determined by minimizing the log ratio of modeled and measured counts over all pairs of tissues between patients (*k≠k*’):

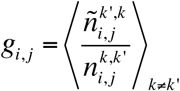

see Fig. S 13 for results and detailed comparison of data to the model.

### Tumor-Normal Classifier

Receiver operator curves (ROC) were computed with the ROCR package in R using the density of T-cells, fraction of unique clonotypes, and clonality (Fig. 1) as metrics to distinguish repertoires from tumor and normal tissue. A classifier based on the coefficient of determination (R^2^) comparing the top 100 clones in blood with tumor and normal breast was also evaluated. Linear regression was performed between the abundances of the top 100 clonotypes in blood and the corresponding abundances in tumor and the fraction of variance captured by the fit (R^2^) tabulated for each sample (Fig. S10). The same process was repeated between the top 100 clonotypes in blood with normal (Fig. S11).

### Enriched Sequences

Sequences between pairs of tissues are plotted on log-log plots where sequences with zero counts in one of the samples are plotted at ¼ of the lowest abundance in that sample. Tumor sequence *n* is enriched if it has abundance greater than 0.1% and relative abundance in the normal exceeding 32, i.e., *T_n_/N_n_* ≥ 32 and including clonotypes not detected in normal breast (i.e., *N_n_* = 0). Similarly, enriched normal sequences are defined as sequences with greater than 0.1% abundance in the normal breast and a ratio with the tumor exceeding 32, i.e., *N_n_/T_n_ ≥ 32*.

### Inter-Patient Sequence Sharing

The number of patients with at least one template detected for each CDR3 sequence is independently tallied for the three tissue compartments and the sequences binned according to the number of shared patients (Fig. S 14). In order to more easily compare sharing between samples and tissues, this process is repeated for the top 1945 CDR3s from each patient (only BR18N has fewer sequences). The expected amount of sharing (dashed lines in Fig. 6A and Fig. S14) is determined by random sampling sequences from a population of size *M*=2×10^6^ according to the sampling depth in each patient and tallying the number of common clonotypes across patients. The median abundance, CDR3 length and number of inserted/deleted nucleotide bases in the junction are computed for each bin. For sequences where the D-gene could not be resolved, the number of deleted bases is conservatively estimated at ten deleted nucleotides, which is the maximum number of deleted D-gene bases reported. Edit distances were calculated using CDR3s with length 13 amino acids from shared sequences (detected in more than one patient) compared with non-shared (only detected in one patient).

### CDR3 Recombination Model

Commonly shared low-diversity sequences are generated from the IMGT germ-line amino acid sequences for the human V-, D- and J-genes. Amino acid sequences contributing to the CDR3 region are extracted from the 5’ of each V-gene starting with the conserved cysteine (or nearest equivalent in some V-genes), the 3’ end of each J-gene ending in the conserved phenylalanine and the full D-gene, including all 3 reading frames. All combinations of these sequences are then used to generate a list of low-diversity CDR3 sequences. To reflect our observation that shared CDR3s are shorter with many deletions and few insertions, all possible amino acid truncations of the D genes and 1-2 amino acids truncations from the 3’ end of the J gene are also included. No inserted amino acids are included. In general, nucleic acid sequences are ignored except when either end of the D-gene or the 3’-end of the J-gene contains 2 nucleic acids thus strongly biasing the amino acid usage at this position. These additional amino acids that are not strictly germ-line but are also included. See Table S 3 for a list of sequences from each gene.

### Database of healthy donors

In order to query the database of 585 healthy bone marrow donors(18), a list of the top 400 shared and 858 enriched CDR3 sequences was compiled. Shared sequences include CDR3s detected in 2 or more tumors or normal breast patients and 4 or more blood samples. These sequences were provided to Adaptive Biotechnologies where an SQL query on their internal database was performed and the subset of ImmunoSeq data from each patient’s matching clonotypes provided to us. For each TCRB sequence the number of patients, the relative abundances, and the corresponding nucleic acid sequences were tallied for enriched/shared sequences and male/female donors.

## Acknowledgements

The authors acknowledge Yasemin Sucu for help in processing samples, Dr. David Hamm and his team at Adaptive Biotechnologies for assistance in the database query, and Florian Rubelt for helpful discussions. The authors are grateful to funding from the Howard Hughes Medical Institute (HHMI), Mattias Westman gift, the John and Marva Warnock Research Fund and the Debra and Andrew Rachleff Research Fund.

## Data availability

All immunosequencing data underlying this study can be analyzed and freely downloaded from the Adaptive Biotechnologies immuneACCESS site at <<TBD>>.

